# The dynamic properties of a nuclear coactivator binding domain are evolutionarily conserved

**DOI:** 10.1101/2021.06.10.447870

**Authors:** Elin Karlsson, Frieda A. Sorgenfrei, Eva Andersson, Jakob Dogan, Per Jemth, Celestine N. Chi

**Affiliations:** Department of Medical Biochemistry and Microbiology, Uppsala University, BMC Box 582, SE-75123 Uppsala, Sweden; Department of Pharmaceutical Biosciences, Uppsala University, BMC Box 582, SE-75123 Uppsala, Sweden

## Abstract

Evolution of proteins is constrained by their structure and function. While there is a consensus that the plasticity of intrinsically disordered proteins relaxes the structural constraints on evolution there is a paucity of data on the molecular details of these processes. The Nuclear co-activator binding domain (NCBD) from CREB-binding protein is a protein-protein interaction domain, which contains a hydrophobic core but is not behaving as a typical globular domain, and has been desribed as ‘molten-globule like’. The highly dynamic properties of NCBD makes it an interesting model system for evolutionary structure-function investigation of intrinsically disordered proteins. We have here compared the structure and biophysical properties of an ancient version of NCBD present in a bilaterian animal ancestor living around 600 million years ago with extant human NCBD. Using a combination of NMR spectroscopy, circular dichroism and kinetic methods we show that although NCBD has increased its thermodynamic stability, it has retained its dynamic biophysical properties in the ligand-free state in the evolutionary lineage leading from the last common bilaterian ancestor to humans. Our findings suggest that the dynamic properties of NCBD have been maintained by purifying selection and thus are important for its function, which includes mediating several distinct protein-protein interactions.

## Introduction

Evolution has shaped proteins into a wide spectrum of structure, stability and dynamics, with fully disordered proteins at one end of the scale and well-folded globular proteins at the other one (*1*). It is becoming clear that dynamic properties *per se* are important for protein function (*1–3*) but it is not trivial to prove that the dynamics are essential for biological function and not merely a general property of proteins. The Nuclear Coactivator Binding Domain (NCBD) is a small (approximately 50 residues) domain from CREB-binding protein (CBP, also called CREBBP), which is a transcriptional coactivator with histone acetylase activity and present in all animals (*4*). The NCBD domain has a hydrophobic core, but is very dynamic and even verging on being intrinsically disordered (*5–7*). It has therefore been described as “molten-globule-like” (*8–11*). The fact that NCBD possesses properties of both intrinsically disordered and globular proteins makes it an interesting system for assessing the role of dynamics in protein structure and how it modulates function. We have previously subjected NCBD to “evolutionary biochemistry” (*12, 13*). Thus, we resurrected and characterized ancestral versions of NCBD and its protein ligand CBP-Interacting Domain (CID) from the NCOA1, 2 and 3 protein family (with three members in human called Src1, Tif2 and ACTR, respectively) to address the evolution of affinity in this protein-protein interaction. The oldest “maximum likelihood” (ML) NCBD variant we could resurrect was present before the Cambrian period, some 540-600 million years ago, in the common ancestor of all present-day animals with bilaterian symmetry. These animals are divided into deuterostomes and protostomes (D/P) and the ancestral NCBD domain is denoted NCBD_D/P_^ML^. It was shown to bind the CID domain with relatively low affinity (*K*_d_ ~ 1-5 μM) whereas more recent variants of NCBD in the vertebrate lineage (ca. 440 million years ago) had acquired present-day affinity for CID (*K*_d_ ~ 0.1-0.2 μM). Structural characterization of ancestral and present-day CID/NCBD complexes showed that the increase in affinity observed for more modern NCBD variants was due to a combination of factors. These include several new interactions driving structural and dynamic changes in the complex, such as formation of a third a helix in CID upon interaction (*13*). In the present paper, we investigate the structural and biophysical properties of the ligand-free state of the ancestral Cambrian-like low-affinity NCBD_D/P_^ML^ and compare it to that of high-affinity present-day NCBD_Human_ to track the evolution of structure and dynamics and their relation to function. The comparison reveals that the overall properties of the ancestral NCBD_D/P_^ML^ domain are very similar to those of the present-day NCBD_Human_. In general, conservation implies function. Thus, this conservation of a highly dynamic structure and molten-globule like properties suggest that these features are subject to purifying selection and hence that dynamics are indeed important for the biological function of NCBD.

## Results

### The structures of ancient NCBD_D/P_ and extant NCBD_Human_ are similar but not identical

Because of its dynamic properties it is challenging to solve the structure of ligand-free NCBD (*8*). First, we performed experiments with ^15^N labeled NCBD_D/P_^ML^. We discovered by serendipity that low pH significantly improved the quality of the ^1^H^15^N heteronuclear single quantum coherence (HSQC) spectrum resulting in well dispersed peaks for NCBD_D/P_^ML^ (Fig. 1), and allowing for a near complete assignment of the backbone and side-chain residues from NMR 3D experiments using ^13^C and ^15^N labeled samples. The NMR structure was thus determined at pH 2.4 by measuring 3D 1H-1H ^13^C/^15^N resolved NOESY experiments for distance restraints determination and ^3^*J*_HNHA_ for dihedral angle determination. The structure of NCBD_D/P_^ML^ was found to be overall consistent with that of the free (*8*) and bound NCBD_Human_ (*6*) and bound NCBD_D/P_^ML^ determined at pH 6.8 (*13*) (Fig. 1). However, a detailed comparison between NCBD_D/P_^ML^ and NCBD_Human_ shows that the orientation of helix 1 (α1) and helix 3 (α3) differ between the structures. Upon structural alignment of NCBD_Human_ and NCBD_D/P_^ML^, we observed that both the α1 and α3 from NCBD_D/P_^ML^ are slightly tilted away relative to that of NCBD_Human_. We also observed that for CID-bound NCBD_D/P_^ML^ and free NCBD_D/P_^ML^ the α3 displayed structural rearrangements (Fig. 2).

**Figure 1.**
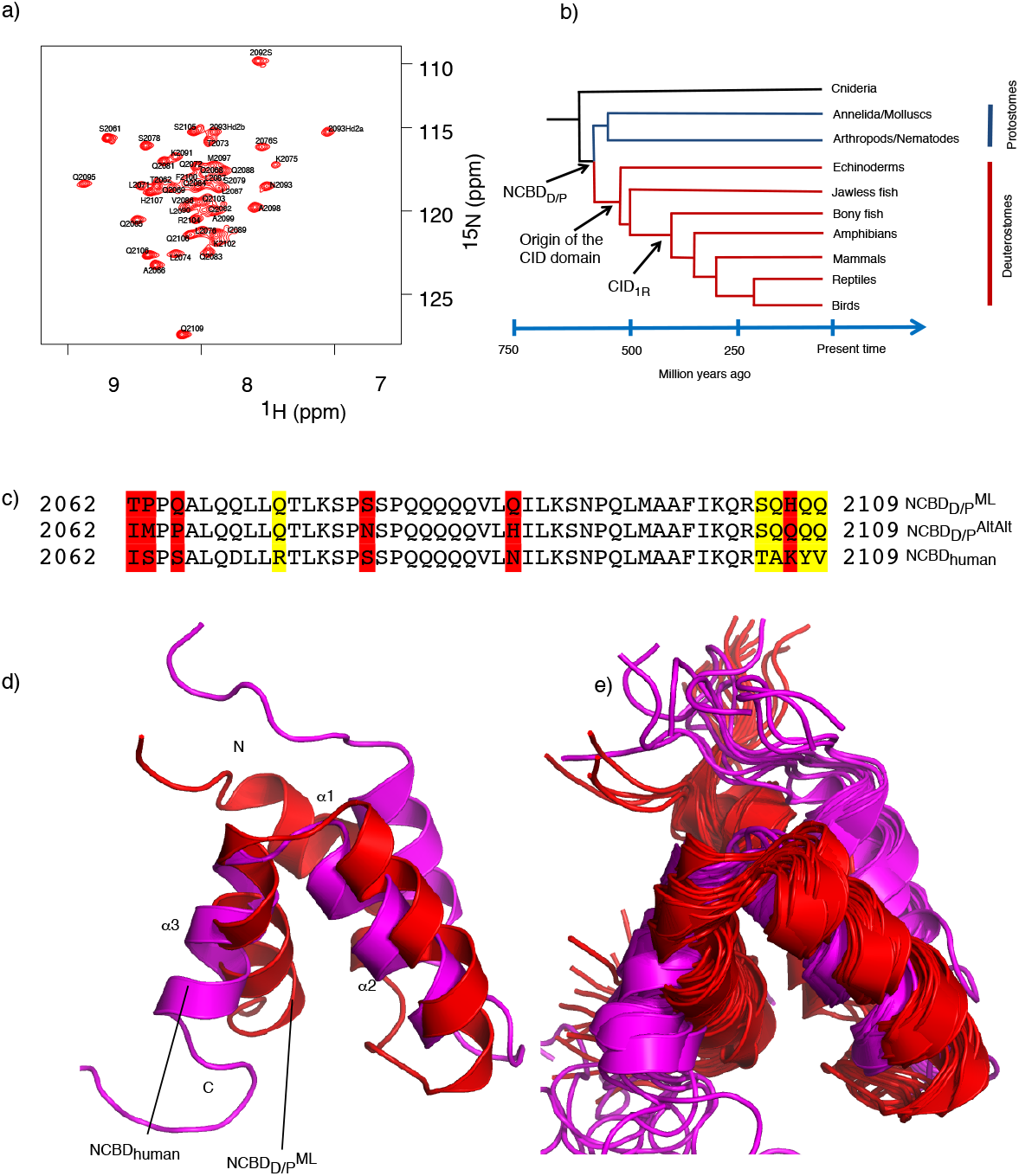
Structures of ancestral and human NCBD domains. (a) ^1^H-^15^N HSQC spectrum of NCBD_D/P_^ML^ at pH 2.4 with assigned residues. All assignments and the structural coordinates have been deposited in the pdb data bank with accession number BMRB ID 34635 and pdb code: 7OSR. (b) A schematic tree depicting the relationship between the deuterostome/protostome (D/P) ancestor and modern species. The times on the x-axis are approximate, in particular the divergence of protostomes and deuterostomes. (c) Sequence alignment for NCBD_Human_, NCBD_D/P_^ML^ NCBD_D/P_^AltAlt^. The residues differing between NCBD_D/P_^ML^ and NCBD_D/P_^AltAlt^ are marked in red while additional differences to NCBD_Human_ are marked in yellow. (d) Comparison of the NMR structure of ancestral NCBD_D/P_^ML^ solved at low pH (red, pdb code: 7OSR), and the previously determined human NCBD (magenta, 2KKJ) (*8*). Note that the structure of human NCBD was solved using a slightly longer construct. (e) Overlay of the 20 lowest energy structures of the NCBD_D/P_^ML^ domain with those of NCBD_human_.

**Figure 2.**
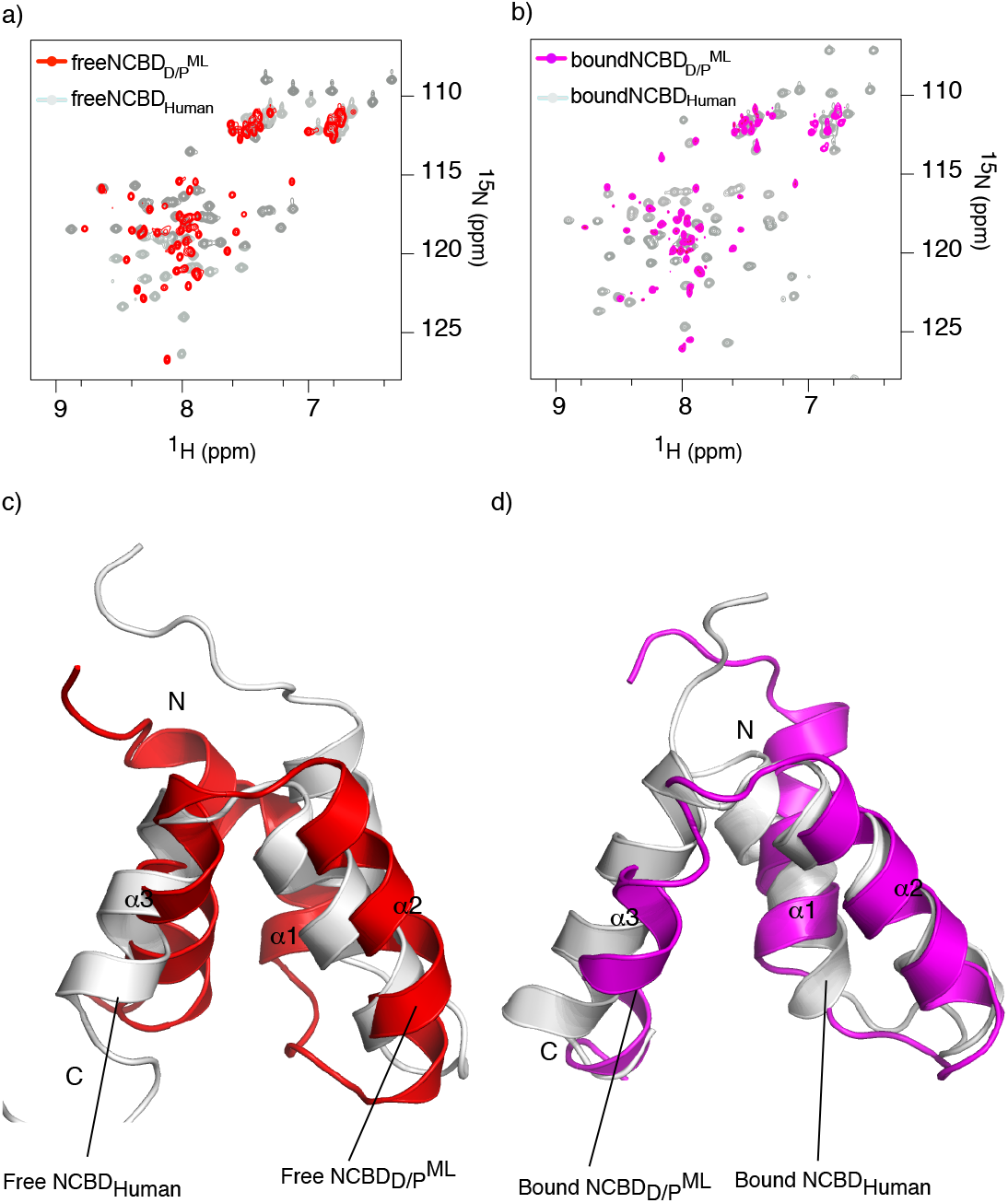
Comparison of the ligand-free and bound ancestral NCBD_D/P_^ML^ and extant NCBD_Human_. The ligand-bound conformations (pdb codes 6ES5 and 6ES7) and the structure of free human NCBD (pdb code: 2KKJ) (*8*) were previously determined. (a) Overlay of the ^1^H-^15^N HSQC spectra between free NCBD_D/P_^ML^ (red) and free NCBD_Human_ (grey). (b) Overlay of the ^1^H-^15^N HSQC spectra between CID-bound NCBD_D/P_^ML^ (magenta) and CID-bound NCBD_Human_ (grey). Structural alignment between (c) free NCBD_D/P_^ML^ (red, 7OSR) and free NCBD_Human_ (grey, 2KKJ), and (d) CID-bound NCBD_D/P_^ML^ (magenta,, 6ES5) and CID-bound NCBD_Human_ (grey, 6ES7).

The amino acid sequence of NCBD_D/P_^ML^ was reconstructed using phylogenetic methods^4^. The probability is low that the resulting maximum likelihood sequence is identical to the actual sequence present in the ancestor. Thus, NCBD_D/P_^ML^ is rather one of a large number of likely ancestral variants with similar properties. We wondered if errors in the sequence of NCBD_D/P_^ML^ resulted in the differences we observe in helix α1 and α3 compared to NCBD_Human_. To investigate this further, and to test how structurally robust the dynamic NCBD domain is, we performed a control experiment where we expressed, purified and characterized an alternative NCBD_D/P_ variant denoted NCBD_D/P_^AltAll^. In this NCBD_D/P_^AltAll^ variant, all residues with a posterior probability lower than around 0.9 (*12*) were replaced with the second most likely residue at that position. For example, residue 2107 is His with 87% probability and Gln with 12% probability in the ancestral NCBD_D/P_. Thus, NCBD_D/P_^ML^ has a His in position 2107 and NCBD_D/P_^AltAll^ a Gln residue. NCBD_D/P_^AltAll^ can be considered a “worst case scenario” variant and a good control of the robustness of conclusions drawn from resurrection experiments. If ML and AltAll variants display similar properties it is very likely that the actual ancestral protein shares these properties as well. Because of this, an AltAll variant is a convenient alternative to combinations of point mutations (*14*). NCBD_D/P_^AltAll^ contains six substitutions as compared to NCBD_D/P_^ML^ (Fig. 1). Three of the differences between NCBD_D/P_^ML^ and NCBD_D/P_^AltAll^ are in the N-terminus, but only one of these is in a structured region, a Gln2065→Pro at the beginning of α1. The other three differences are Ser2078→Asn in the loop between α1 and α2, a solvent exposed Gln2088→His in α2 and His2108→Gln at the C-terminus. We solved the NMR structure of NCBD_D/P_^AltAll^ (Fig. S1) and found that it folds into a similar structure as the other NCBD domains. The overall RMSD between NCBD_D/P_^AltAll^ and NCBD_D/P_^ML^ was 7.9 Å. As a comparison, the overall RMSD between NCBD_Human_ and NCBD_D/P_^AltAll^ or NCBD_D/P_^ML^ were 4.5 Å and 7.0 Å, respectively. Detailed inspection of the structures revealed differences in the backbone between NCBD_D/P_^ML^ and NCBD_D/P_^AltAll^, especially in the orientations of the helices. An overlay of the structures of NCBD_D/P_^AltAll^, NCBD_D/P_^ML^, and NCBD_Human_, shows differences between α1, α2 and α3 (Fig. S1). One reason for the differences is that there are several long-range NOEs in NCBD_D/P_^AltAll^ that are not present in NCBD_D/P_^ML^, between the following pairs of residues: 2068/2082, 2069/2080, 2070/2083 and 2071/2099. There is also an NOE between Ile2062 Hg, and Pro2065 Ha, two of the substituted residues in NCBD_D/P_^AltAll^. However, there appears to be a consistency in the hydrophobic core of both NCBD_D/P_^ML^ and NCBD_D/P_^AltAll^ since the single Phe residue at position 2100 is packed and buried in a similar fashion in the two variants. Phe2100 is stabilized by Leu2090 in NCBD_D/P_^ML^, and both Ile2089 and Leu2090 in NCBD_D/P_^AltAll^ (Fig. 3). Interestingly, this is the same kind of packing that was observed in the free NCBD_human_ domain (*8*), suggesting conserved features in the hydrophobic core.

**Figure 3.**
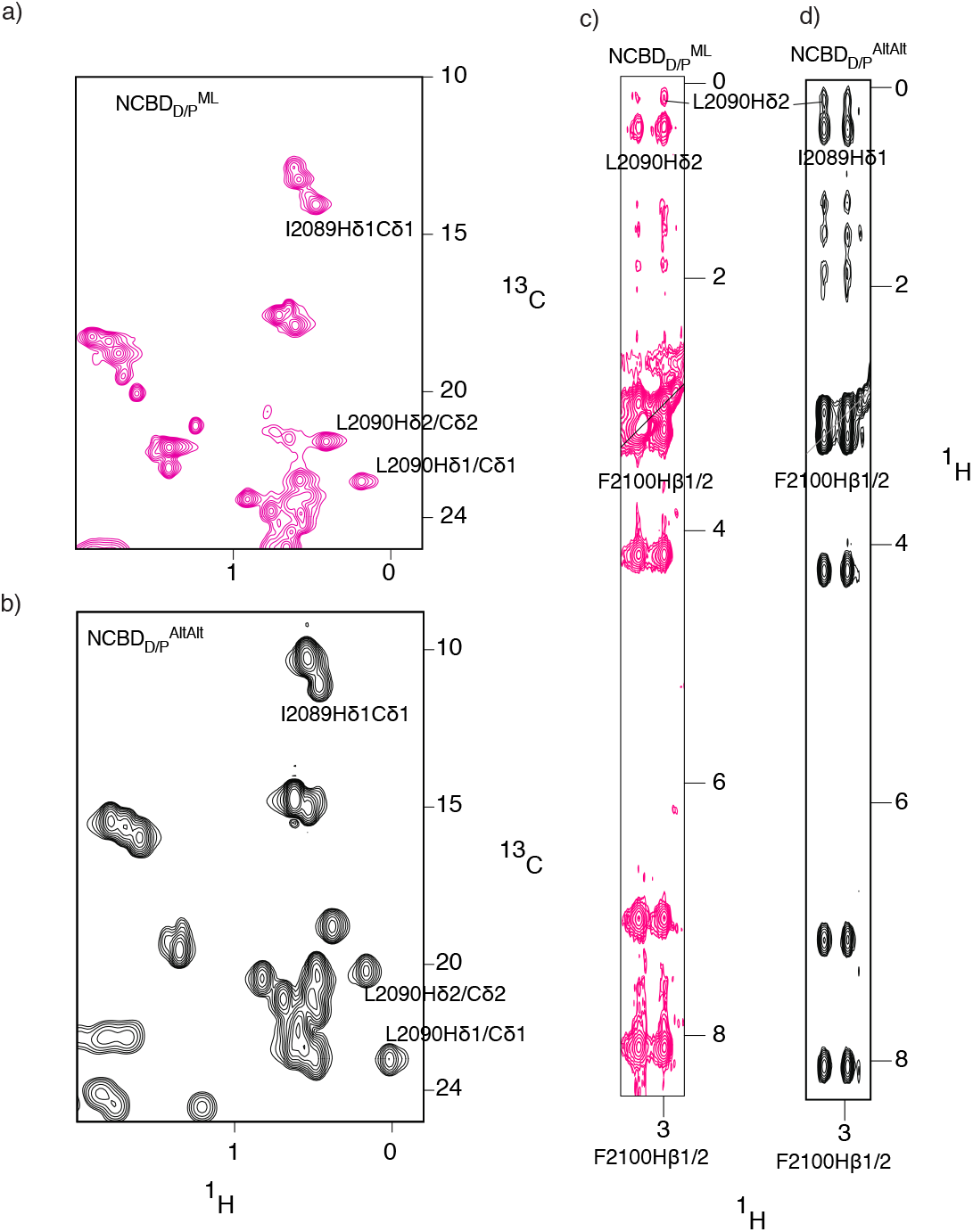
Phe2100 makes NOEs to Ile2089 and Leu2090 in NCBD_D/P_^ML^ and NCBD_D/P_^AltAlt^. (a) ^1^H-^13^C HSQC spectra of the methyl region of NCBD_D/P_^ML^, showing the Hδ1-Cδ1 shifts of L2090 and I2089. (b) ^1^H-^13^C HSQC spectra of the methyl region of NCBD_D/P_^AltAlt^, showing the Hδ1-Cδ1 shifts of L2090 and I2089. (c) Strip from a ^1^H-^1^H ^13^C resolved NOESY showing NOEs from the Hβ1/2 atoms of Phe2100 and the associated NOEs to Leu2090Hδ1 for NCBD_D/P_^ML^. (d) Strip from a ^1^H-^1^H ^13^C resolved NOESY showing NOEs from the Hβ1/2 atoms of Phe2100 and the associated NOEs to Leu2090Hδ1 and Ile2089Hδ1 for NCBD_D/P_^AltAlt^. These signature NOEs implies that both proteins form a hydrophobic core similar to that of NCBD_Human_.

Despite the differences discussed in the previous paragraph, the overall biophysical properties of NCBD_D/P_^ML^ and NCBD_D/P_^AltAll^ were found to be similar (Fig. S2) corroborating the overall similarity seen with the NMR structures. Specifically, to test the affinity of NCBD_D/P_^AltAll^ for CID, we expressed and purified ML and AltAll versions of its ligand CID_1R_ based on our previous reconstruction (*8*). *K*_d_ values of all four combinations (NCBD_D/P_^ML^ and NCBD_D/P_^AltAll^ with CID_1R_^ML^ and CID_1R_^AltAll^) were between 2-9 μM (Fig. S2a), which is in the same range as previous estimates of affinity of the ancestral complex (*12, 13*). Furthermore, the stability as monitored by urea denaturation (Fig. S2b) and thermal denaturation (Fig. S2c-d) were similar for NCBD_D/P_^ML^ and NCBD_D/P_^AltAll^. Thus, based on the highly similar biophysical properties of NCBD_D/P_^ML^ and NCBD_D/P_^AltAll^ we conclude that our conclusions are robust with regard to the uncertainty in the reconstructed sequence.

### Ancient NCBD_D/P_ is less thermodynamically stable than extant NCBD_Human_ but display a similar pH dependence with regard to structure and stability

Since the NCBD_D/P_^ML^ structure was determined at low pH, we next investigated how pH influences the fold and stability. First, we performed ^1^H-^15^N HSQC-monitored pH titrations of NCBD_D/P_^ML^ and compared them to those of NCBD_Human_ at different pH values (Fig. 4). The NMR titration experiments showed that NCBD_D/P_^ML^ is as folded at low pH as it is at high pH values. However, the peaks (^1^H-^15^N) are much sharper (reduced line-width) at lower pH values indicating less conformational heterogeneity. Second, the CD-monitored urea denaturation experiments showed identical stability pattern at two pH values (3.0 and 7.4) for both ancient NCBD_D/P_^ML^ and extant NCBD_Human_ (Fig. 4c and d, Supplementary table S1). This indicates that while at lower pH the NCBD_D/P_^ML^ still exhibits the same stability as at high pH values, there is a shift in conformational heterogeneity towards a single state at low pH. Furthermore, whereas the apparent thermodynamic stability differs by approximately 1 kcal mol^-1^ for ancient NCBD_D/P_^ML^ and extant NCBD_Human_, both are largely unaffected by a drop in pH to 3.0 (Supplementary Table S1). The observed difference in stability between NCBD_D/P_^ML^ and NCBD_Human_ appears to be a robust result. Previous experiments suggested that the ancestral NCBD_D/P_^ML^ was slightly less stable than younger and extant NCBD variants as judged by urea denaturation experiments at pH 7.5 (*12*). However, the broad unfolding transition of NCBD resulting from its small hydrophobic core leads to a large error in the *m*_D-N_ value, which makes it difficult to unequivocally determine the thermodynamic stability of the domain. However, all present results (including experiments on the stabilized NCBD_D/P_^ML^ variant NCBD_D/P_^T2073W^ (Supplementary Table S1) agree well with the previous experiments and we can therefore conclude with greater confidence that the ancestral NCBD_D/P_^ML^ was less thermodynamically stable than evolutionarily younger variants. Interestingly, with some exceptions (*15, 16*), ancestral reconstructions often yield proteins with higher thermostability, which correlates with thermodynamic stability, than extant proteins, either due to a bias for stability in the reconstruction or due to higher temperatures in past times (*17–21*), like during Cambrian (*22, 23*). In any case, ancestral NCBD appears to have populated the native state to a lesser extent than present day human NCBD.

**Figure 4.**
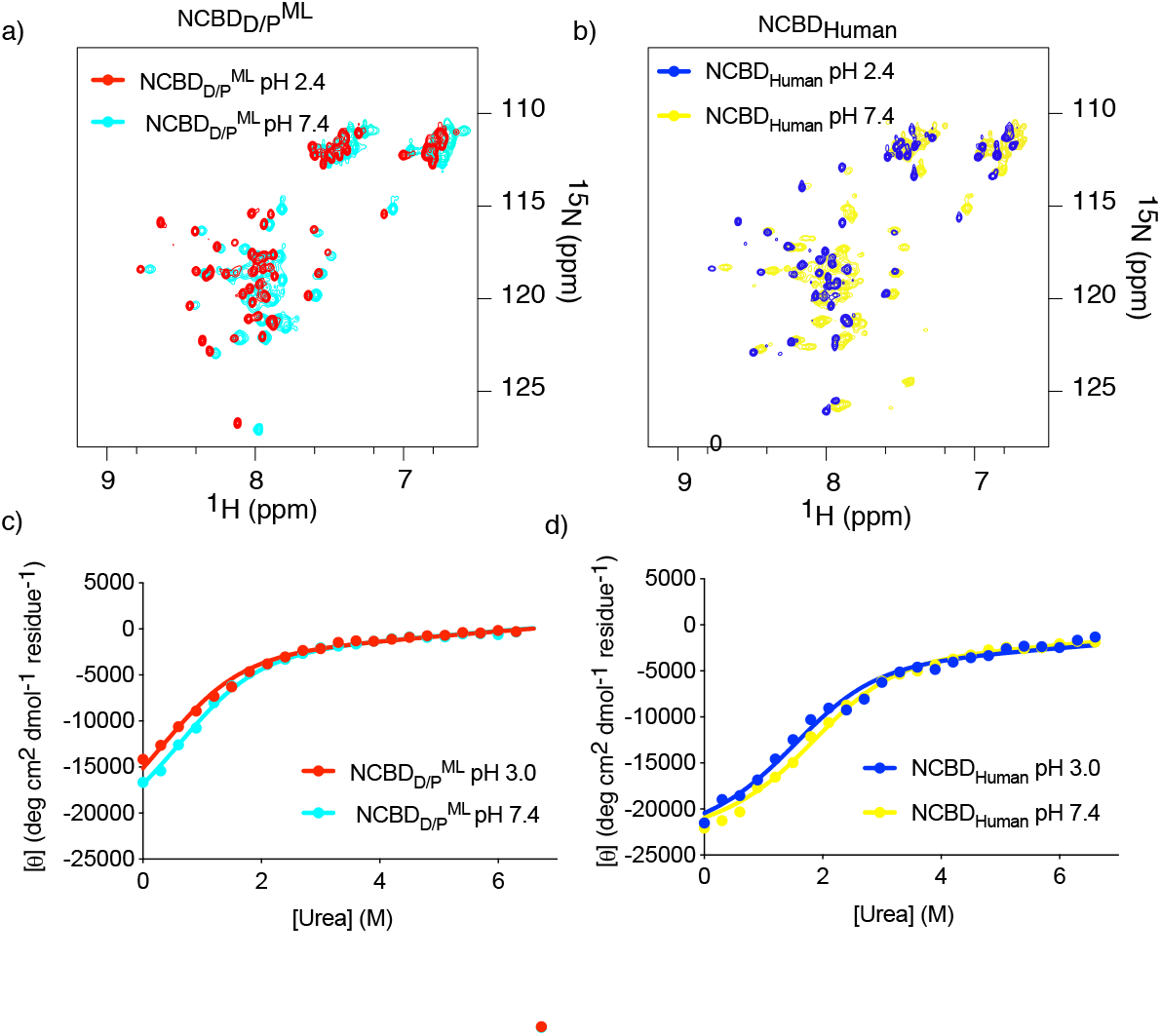
Comparison of ancestral NCBD_D/P_^ML^ and extant NCBD_Human_ at two different pH values. a) Overlay of the ^1^H-^15^N HSQC spectra of NCBD_D/P_^ML^ at pH 2.4 (red) and at pH 6.8 (cyan). b) Overlay of the ^1^H-^15^N HSQC spectra of NCBD_Human_ at pH 2.4 (blue) and at pH 6.8 (yellow). c) Urea denaturation experiments for NCBD_D/P_^ML^ at pH 3.0 (red) and at pH 7.4 (cyan). d) Urea denaturation experiments for NCBD_Human_ at pH 3.0 (yellow) and at pH 7.4 (blue).

### Ancient NCBD_D/P_^ML^ and extant NCBD_Human_ display similar temperature-dependent structural changes

Previous experiments demonstrated that NCBD_Human_ shows an apparent non-cooperative unfolding behavior when subjected to increasing temperatures, as monitored by CD (*6, 8*). This behavior may result from a low Δ*H*_D-N_ value rather than a true non-cooperative unfolding. (The urea denaturation experiments are consistent with a cooperative two-state unfolding.) We compared ^1^H-^15^N HSQC and CD spectra, respectively, of NCBD_D/P_^ML^ and NCBD_Human_ at different temperatures (Fig. 5). In agreement with the previous data we observed that both variants unfold non-cooperatively with temperature as monitored by the increase in molar ellipticity at 222 nm suggesting loss of a-helix as the temperature is increased. However, for both NCBD_D/P_^ML^ and NCBD_Human_ the signal is not completely lost even at 363 K (90°C) underscoring that this is not a typical native to highly-disordered-denatured state transition. Furthermore, ^1^H-^15^N HSQC spectra of NCBD_D/P_^ML^ at temperatures up to 333 K (60°C) show well dispersed peaks indicating a presumably globular, collapsed architecture (Fig. S3) and corroborating the notion that NCBD retains structure at high temperatures.

**Figure 5.**
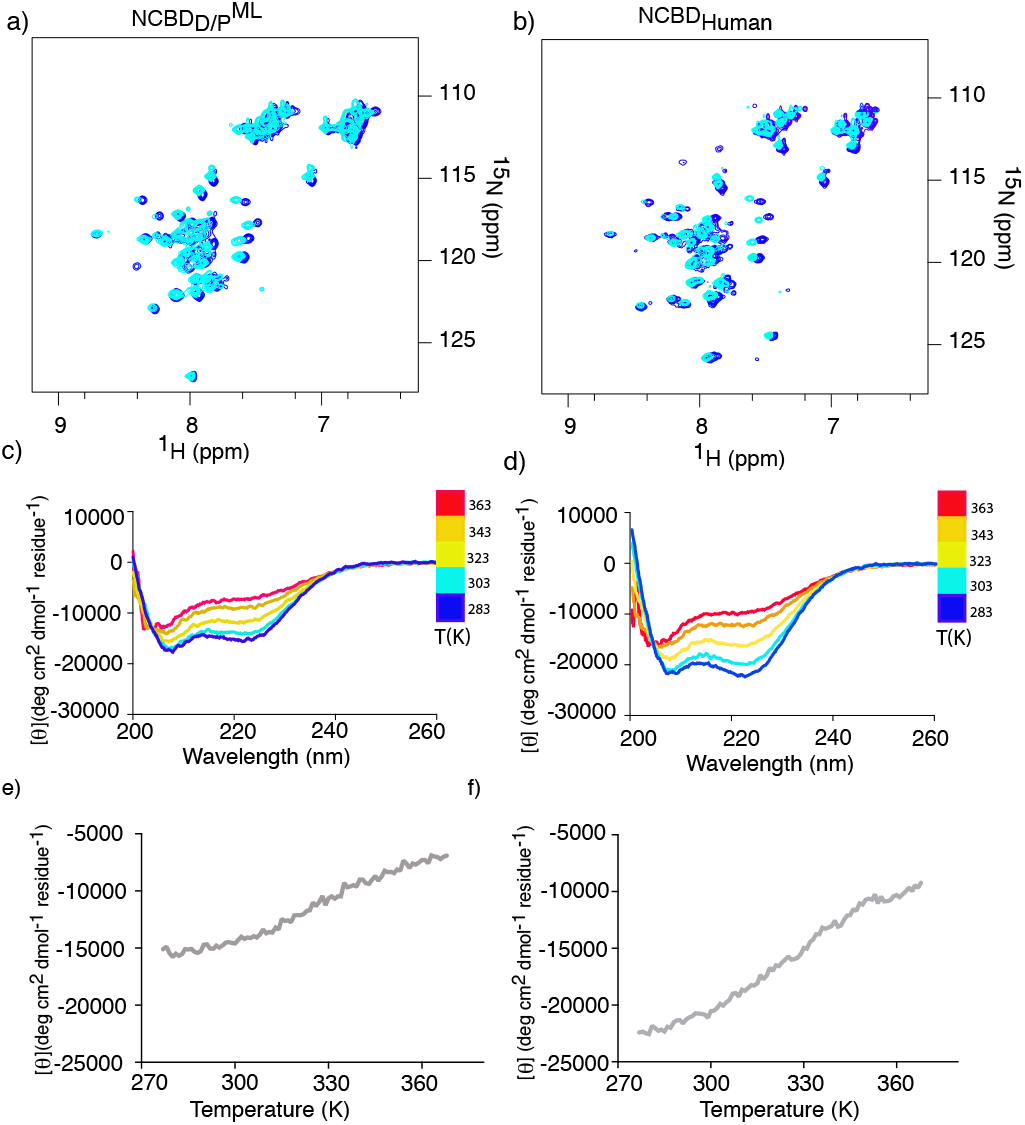
Temperature dependence for ancestral NCBD_D/P_^ML^ and extant human NCBD. a) Overlay of ^1^H-^15^N HSQC spectra of NCBD_D/P_^ML^ at 283 K (blue) and at 303 K (cyan). b) Overlay of ^1^H-^15^N HSQC spectra of NCBD_Human_ at 283 K (blue) and at 303 K (cyan). Circular dichroism spectra at a range of temperatures (283-363 K) for (c) NCBD_D/P_^ML^ and (d) NCBD_Human_. Molecular ellipticity at 222 nm (probing a-helix content) as a function of temperature for (e) NCBD_D/P_^ML^ and (f) NCBD_Human_.

### Rate constants for (un)folding of ancient NCBD_D/P_^ML^ and extant NCBD_Human_

The folding of human NCBD is complex, and, based on kinetic experiments on a human NCBD variant, we proposed a three-state system involving ‘native’ NCBD (*i.e*., the solved structure), an intermediate state with unknown structure and a denatured state, which likely retains significant structure under native conditions (*24*). Rate constants in the same order was determined by relaxation dispersion NMR experiments by Kjaergaard et al. (*10*) It is clear from the NMR experiments that low pH promotes the native state of NCBD and that pH does not affect the thermodynamic stability as monitored by urea denaturation experiments to any significant extent, neither for NCBD_D/P_^ML^ nor NCBD_Human_. Previous folding experiments on NCBD_Human_ (*24*) involving jumps from low to neutral pH thus resulted in small shifts in the equilibria between the native, intermediate and denatured states yielding the observed kinetic transients. In fact, a likely explanation for the better NMR spectra at low pH is that there is less of the intermediate state under these conditions.

To further compare the biophysical properties of ancient NCBD_D/P_^ML^ and extant NCBD_Human_ we performed kinetic folding experiments using stopped-flow as well as temperature-jump fluorescence spectroscopy. To achieve this we introduced a Trp at position 2073 in NCBD_D/P_^ML^ as previously done in the folding studies of NCBD_Human_ (*24*). This modification resulted in a thermodynamic stabilization of around 1 kcal mol^-1^ for both NCBD_D/P_^T2073W^ and NCBD_Human_^T2073W^ (Fig. 6a-b, Supplementary Table S1). Previous folding experiments were performed on a slightly longer NCBD_Human_ variant (residues 2058-2116), which was used in the original studies of CID/NCBD (*6*). In our resurrection studies, we used a shorter version of NCBD_Human_, only containing the evolutionarily conserved region (residues 2062-2109). Therefore, we first repeated the temperature-jump folding experiments for NCBD_Human_^T2073W^ and showed that the shorter construct used in our evolutionary studies displayed similar kinetics as the longer construct (Fig. 6c). We then performed folding experiments with NCBD_D/P_^T2073W^. The kinetic transients of NCBD_D/P_^T2073W^ from temperature jump experiments had lower signal-to-noise than those of NCBD_Human_^T2073W^, but observed rate constants were in the same range (Fig. 6d). To investigate the folding kinetics further we used stopped-flow fluorimetry, which has a lower time resolution but can be used to compare folding rate constants for NCBD_D/P_^T2073W^ and NCBD_Human_^T2073W^ at low temperature (Fig. 6e-f). The stopped-flow folding experiments were conducted by making pH jumps from a 5 mM HCl solution (pH ~ 2) to a buffer solution of 20 mM sodium phosphate (pH 7.4, 150 mM NaCl, 1 M TMAO) and monitoring the relaxation through a 330 nm band-pass filter at 4°C. The low temperature is necessary to reduce the observed rate constant such that the folding or other conformational transitions take place in the millisecond time window. We observed two kinetic phases for NCBD_Human_^T2073W^ (one fast phase with a negative amplitude and a second slower phase with a positive amplitude) but only one kinetic phase for NCBD_D/P_^T2073W^ (with a positive amplitude). The values of the observed rate constants for NCBD_Human_^T2073W^ (~300 s^-1^ and ~70 s^-1^, respectively) were similar to those observed previously for the longer construct (*24*). The single value obtained for the NCBD_D/P_^T2073W^ was ~140 s^-1^. Trp fluorescence is a very sensitive but crude structural probe. Thus, the number of kinetic phases in folding experiments could be dependent on small structural rearrangements around the Trp. Nevertheless, the observed rate constant for NCBD_D/P_^T2073W^ in the stopped flow pH jump experiment is in the same range as the two rate constants observed under the same conditions for human NCBD.

**Figure 6.**
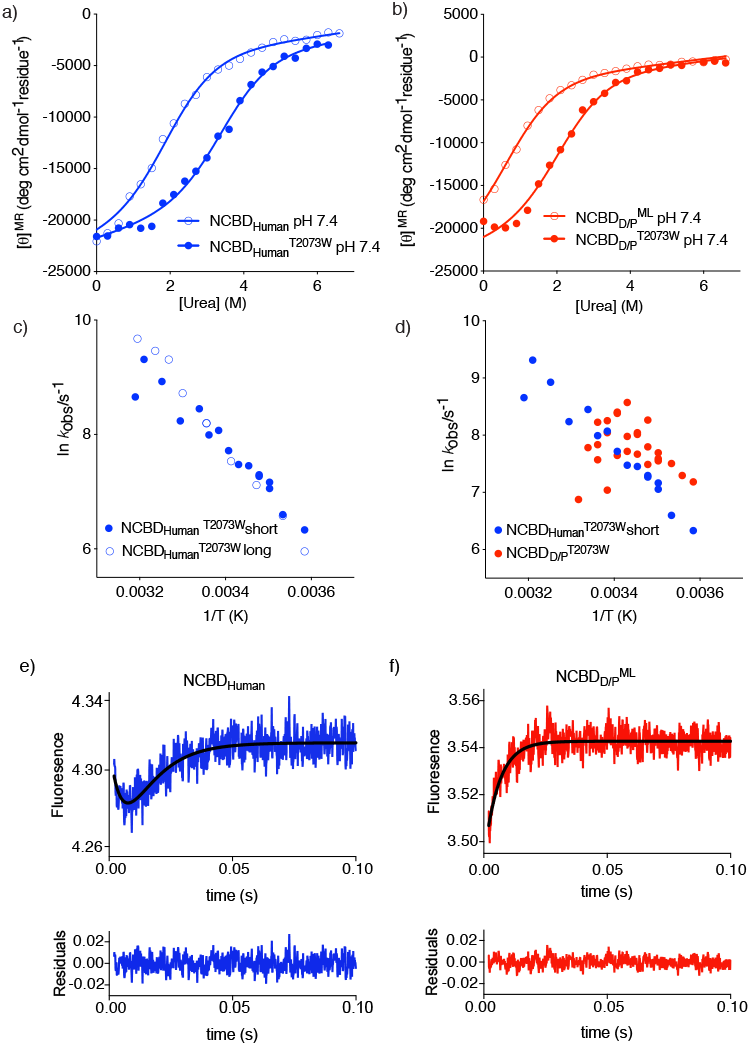
Stability and temperature dependence of kinetic rate constants for NCBD_D/P_^ML^ and extant human NCBD. (a) Stability measurement of NCBD_Human_^WT^ (open circles) and NCBD_Human_^T2073W^ (filled circles) measured by urea denaturation. (b) Stability measurement of NCBD_D/P_^ML^ (open circles) and NCBD_D/P_^T2073W^ (filled circles). Parameters from the curve fitting are shown in Supplementary Table S1. (c) Temperature jump kinetic experiments for a longer human NCBD construct used in previous studies (*24*) (open circles) and the shorter human NCBD variant used in this study (filled circles). Both variants contained the T2073W mutation. (d) A comparison of observed rate constants from temperature-jump experiments for NCBD_Human_^T2073W^ (blue) and NCBD_D/P_^T2073W^ (red). All experiments were performed in 20 mM sodium phosphate pH 7.4, 150 mM NaCl. (e-f) Kinetic (un)folding of NCBD_Human_^T2073W^ and NCBD_D/P_^T2073W^ measured by stopped flow fluorescence spectroscopy. (e) (Un)folding kinetic transient of NCBD_Human_^T2073W^ (blue) fitted to a double exponential function (black line). Below each transient are the residuals from the respective fits. (f) Stopped flow (un)folding kinetic transient of NCBD_D/P_^T2073W^ (red) fitted to a single exponential function (black line). The experiments were performed by jumping from 5 mM HCl (pH ~ 2) to a final buffer concentration of 20 mM sodium phosphate (pH 7.4), 150 mM NaCl, 1 M TMAO. The relaxation was monitored using a 330 band-pass emission filter. Fitting a double exponential function resulted in *k*_obs_ values of 290 ± 30 and 67 ± 4 s^-1^ for NCBD_Human_^T2073W^. Fitting a single exponential function yielded a *k*_obs_ value of 139 ± 7 s^-1^ for NCBD_D/P_^T2073W^.

## Discussion

The biophysics and molecular evolution of NCBD is interesting for several reasons, which all relate to the dynamic properties of NCBD. Firstly, while it has a hydrophobic core and globular shape, NCBD is a very dynamic protein domain with characteristics of an IDP (*5, 6, 8–11, 25*). Secondly, NCBD has several binding partners including the three NCOA transcriptional co-regulator paralogs (*5, 26*), the transcription factors p53 (*27*), Ets-2 (*28*) and interferon factor-3 (*29*), and viral proteins such as Tax (*30*) from T-cell leukemia virus. Intriguingly, NCBD displays great conformational plasticity as shown by its complex with interferon factor-3 where the helices of NCBD adopt a distinct conformation as compared to the complex with CID (*31*). Thirdly, evolutionary snapshots of the complex between CID and NCBD suggest that several new intermolecular contacts and conformational plasticity increased the affinity when going from the Cambrian-like NCBD_D/P_^ML^ (*K*_d_ ~ 1-5 μM) to younger variants (*K*_d_ ~ 0.2 μM) (*12, 13, 32*). Fourthly, while affinity and structural order of modern NCBD/CID complexes were increased, the transition state of the folding-induced binding displayed more conformational heterogeneity as compared to the ancestral complex (*33*). Our present results, how the free, unbound state of NCBD has evolved during the transition from a low-affinity to a high-affinity CID binder relate to all four points.

The most striking finding is that NCBD_D/P_^ML^ displays very similar molten-globule like properties as the previously characterized human NCBD (*8, 10*). This conservation of properties, borderline between a globular domain and an IDP, suggests that the dynamics of NCBD is preserved by purifying selection and therefore of functional importance. In general, structural plasticity is regarded as one main functional feature of IDPs allowing for multiple protein-protein interactions (*34*). NCBD interacts with several protein domains, both folded and disordered ones, and the dynamic nature of NCBD likely increases its propensity to adapt to different partners. Moreover, NCBD interactions result in coupled folding and binding. Many such reactions have recently been shown to occur via a mechanism denoted templated folding, where the binding partner influences the folding pathway (*35–37*). Indeed, this was observed for the interaction between human NCBD and CID (*33, 38, 39*). Thus, our present findings suggest that templated folding may be a general feature of binding reactions with NCBD where the conserved dynamics facilitate sampling of different free states, heterogeneity of bound states, as well as the pathway between free and bound states.

Going into details of the NMR model, we note that while overall structure is robust to errors in the predicted sequence as shown by the comparison of the NMR models of free NCBD_D/P_^ML^ and NCBD_D/P_^AltAll^, there are differences in for example helix orientations. Globular domains with a sequence identity as high as these NCBD variants (77-88%) would typically have identical structures due to a highly funneled energy landscape (*40, 41*), while proteins with larger structural heterogeneity have less funneled landscapes (*1*). Indeed, the differences between the structures of NCBD_Human_, NCBD_D/P_^ML^ and NCBD_D/P_^AltAll^, in particular regarding the angles of the helices in relation to each other, are consistent with a less funneled energy landscape resulting in conformational heterogeneity (*11*). This heterogeneity is also consistent with the conserved dynamic properties of NCBD. In conclusion, our data suggest evolutionarily conserved dynamics of NCBD that underlie functional plasticity in ground state conformations, templated binding-and-folding and a propensity to interact with several binding partners.

## Materials and methods

### Protein sequences, expression and purification

The reconstruction of ancestral sequences was previously published (*12*). The AltAll versions of NCBD_D/P_ (Fig. 1) and CID 1R used in the present paper was based on posterior probabilities from the reconstruction with a cutoff of 91%. One position in NCBD_D/P_ (Thr2062) is very uncertain. A Thr residue (15%) was originally chosen at this position in NCBD_D/P_^ML^ to avoid a hydrophobic residue, although Ile (19%) has a higher posterior probability. In the current NCBD_D/P_^AltAll^ residue 2062 is Ile. NCBD and CID variants were expressed in *E. coli* from a pRSETA plasmid and purified as previously described (*33*) Concentrations were determined by absorbance at 280 nm for NCBD (using calculated extinction coefficients based on amino acid composition) or 205 nm for CID using an extinction coefficient of 250,000 M^-1^cm^-1^. Purity was checked by SDS-PAGE and identity of the purified proteins was confirmed by MALDI-TOF mass spectrometry.

### Circular dichroism

Far-UV CD experiments were performed using a JASCO J-1500 CD spectrophotometer with a Peltier temperature control system. The protein concentration in all CD experiments was 18-20 μM. The buffer solution was either 20 mM sodium phosphate (pH 7.4), 150 mM NaCl or 50 mM potassium formate (pH 3.0), 150 mM NaCl, and CD spectra were recorded at 277 K. The scanning speed was 50 nm/min, bandpass 1 nm and integration time 1 second. Each reported spectrum is an average of 3-4 recorded spectra. In the equilibrium thermal denaturation experiments the proteins were dissolved in 50 mM potassium formate buffer (pH 3.0), 150 mM NaCl and the sample was heated from 277 to 368 K. The heating rate was 1 K/min, waiting time 5 seconds and the denaturation was monitored at 222 nm. During the heating, spectra (average of three measurements) were taken at 283, 303, 323, 343 and 363 K, respectively. In the equilibrium urea denaturation experiments the proteins were denatured in 0-6.6 M urea at 277 K and the unfolding was monitored at 222 nm. The data was analyzed in GraphPad prism and fitted by non-linear regression to a two-state equilibrium model (*42*). Because of the broad unfolding transition and lack of native baseline for several variants, four parameters were shared among all urea denaturation data sets in the curve fitting (Fig. S4, Supplementary Table 1). One of these parameters was the molar ellipticity of the native state, with the reasonable assumption that the secondary structures of the native states are similar for all variants. The other parameters were the slopes of the denatured and native baselines as well as the m_D-N_ value (assuming a similar change in solvent accessible surface area upon denaturation for all variants).

### Temperature jump experiments

Temperature jump experiments were conducted using 100-200 μM protein on a TJ-64 temperature jump system (TgK Scientific). The buffer solution was 20 mM sodium phosphate (pH 7.4), 150 mM NaCl. The proteins were subjected to temperature jumps of 2-8.5 K to different target temperatures. The data were fitted in the Kinetic Studio software (TgK scientific) to a double exponential function, to account both for the fast phase (relaxation time ~ 80 μs), which corresponds to the heating time of the instrument, and to the slow phase, which corresponds to the conformational transition in the protein. A Trp variant of NCBD_D/P_^ML^ (NCBD_D/P_^T2073W^) was chosen for kinetic experiments since a Trp in this position was previously used for human NCBD. We engineered a Trp at other positions in NCBD_D/P_^ML^ and obtained rate constants in the same range for all variants (Supplementary Table S2).

### Stopped flow experiments

The stopped flow experiments were performed using an upgraded SX-17MV stopped flow spectrophotometer (Applied Photophysics) and the measurements were conducted at 277 K. Proteins were dissolved in 5 mM HCl (pH ~ 2) and rapidly mixed 1:1 with buffer solutions complemented with trimethylamine N-oxide (TMAO) to a final concentration of 20 mM sodium phosphate (pH 7.4), 150 mM NaCl, 1 M TMAO. The final protein concentration was 5 μM. The excitation wavelength was 280 nm and the resulting fluorescence emission was detected after passage through a 330 nm bandpass filter. The transients were analyzed in GraphPad Prism and fitted to a single or double exponential function.

### NMR experiments

NMR experiments were acquired on Bruker 600, 700 and 900 MHz spectrometers equipped with triple resonance cryogenic temperature probes at 298 K except otherwise stated. The final NMR samples contained 500 μM protein to which 0.01% NaN_3_ and 5% D_2_O was added. Experiments for assignment and subsequent structure determination were done at pH 2.4 (HCl adjusted with NaOH) for NCBD_D/P_^ML^ and pH 6.8 for NCBD_D/P_^AltAll^. The following NMR experiments were recorded for assignment and subsequent structure determination: standard 2D ^1^H-^15^N HSQC, 3D HNCACB ^15^N-resolved [^1^H-^1^H]-NOESY-HSQC, ^15^N-TOCSY-HSQC ^13^C-resolved [^1^H-^1^H]-NOESY-HSQC, HCCH-TOCSY and ^13^C-resolved [^1^H-^1^H]-TOCSY-HSQC. Phi-angle restraining ^3^*J*_HNHA_ couplings were determined from 3D HNHA type experiments using quantitative *-J* intensity modulated experiments (*43*). The temperature variation experiments were performed in 20 mM sodium formate, pH 3.0. All experiments were processed with nmrpipe (*44*) and analysed with CCPnmr (*45*).

### Structure calculation

Structure calculations were done using the CYANA 3.97 (*46*) package as follows: Initially, cross peaks were converted into upper distance restraints following an automated process in CYANA. These distance restraints together with *φ/ψ* dihedral angles determined from C^α^-chemical shifts and ^3^*J*_HNHα_ (measured) were used as input for the initial structure calculations. The structures were calculated with 200,000 torsion angle dynamics steps for 100 conformers starting from random torsion angles by simulated annealing. 20 conformers with the lowest target function values were selected and analyzed. The structural statistics together with all input data for the structure calculations are presented in Supplementary Table S3. Assignments and structural coordinates have been deposited in the RCSB protein data bank with pdb code: 7OSR (accession number BMRB ID 34635) (NCBD_D/P_^ML^) and 7OSW (accession number BMRB ID 34636) (NCBD_D/P_^AltAll^).

## Acknowledgements

This work was supported by the Wenner-Gren foundation WG-17 returning grants to C. N. C., and by the Swedish Research Council (2020-04395 to P.J.) and Knut and Alice Wallenberg foundation (Evolution of new genes and proteins, to P.J.). We used the NMR Uppsala infrastructure, which is funded by the Department of Chemistry - BMC and the Disciplinary Domain of Medicine and Pharmacy. We thank Magnus Kjaergaard for valuable comments on the NMR experiments.

## Supplementary information

### Supplementary figures

**Figure S1.**
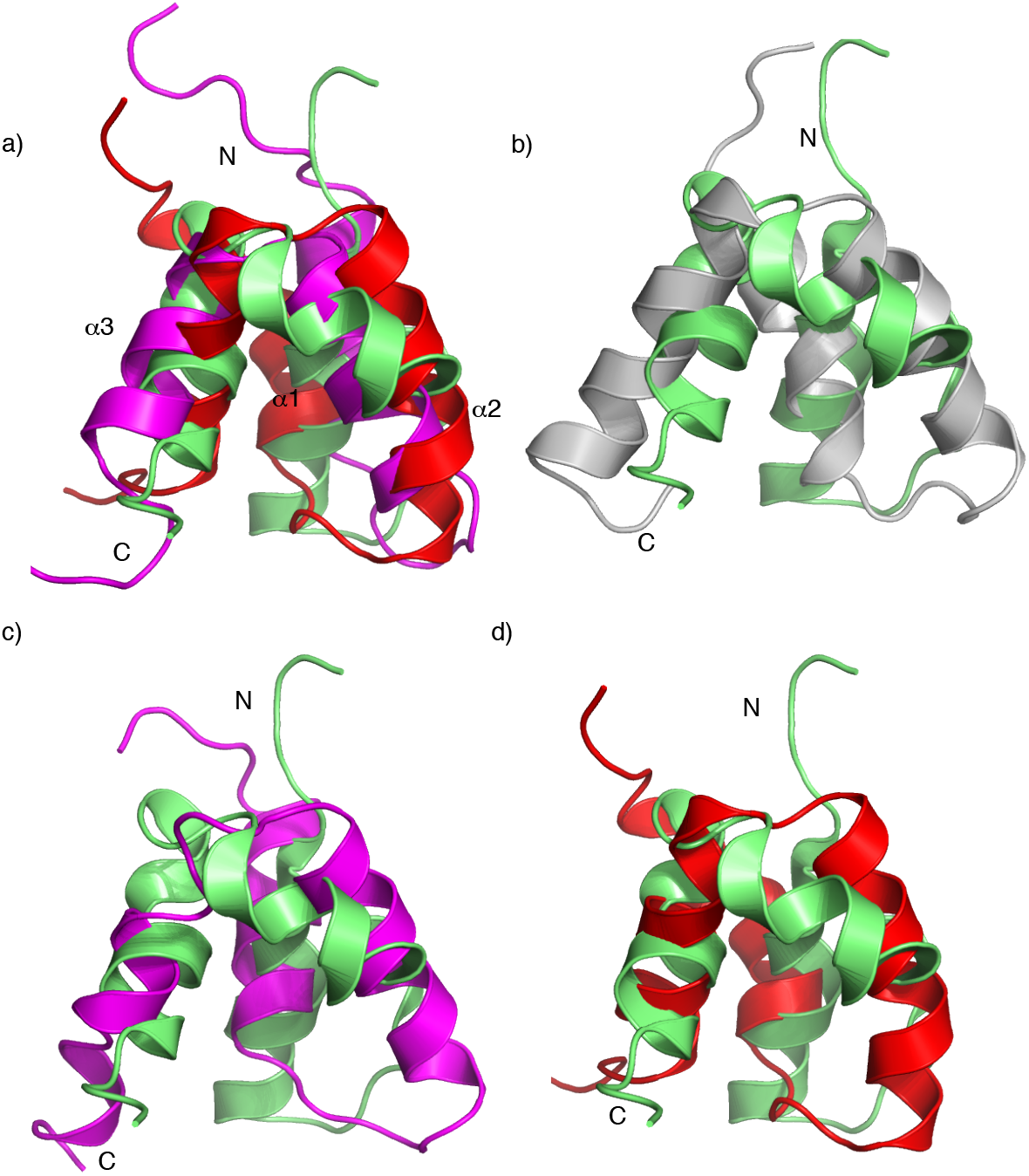
Overlay of the structures of NCBD_D/P_^ML^, NCBD_D/P_^AltAlt^ and NCBD_Human_. (a) An overlay of the structures determined from NMR for NCBD_D/P_^ML^(red, pdb code: 7OSR), NCBD_D/P_^AltAlt^ (green, 7OSW), and the previously determined NCBD_Human_ (magenta, 2KKJ). (b) Overlay of NCBD_D/P_^AltAlt^ (green, 7OSW), and the previously determined CID-bound NCBD_D/P_^ML^ (grey, 6ES5). (c) Overlay of NCBD_D/P_^AltAlt^ (green, 7OSW), and the previously determined NCBD_Human_ (magenta, 2KKJ). (d) Overlay of NCBD_D/P_^AltAlt^ (green, 7OSW) and NCBD_D/P_^ML^ (red, 7OSR).

**Figure S2.**
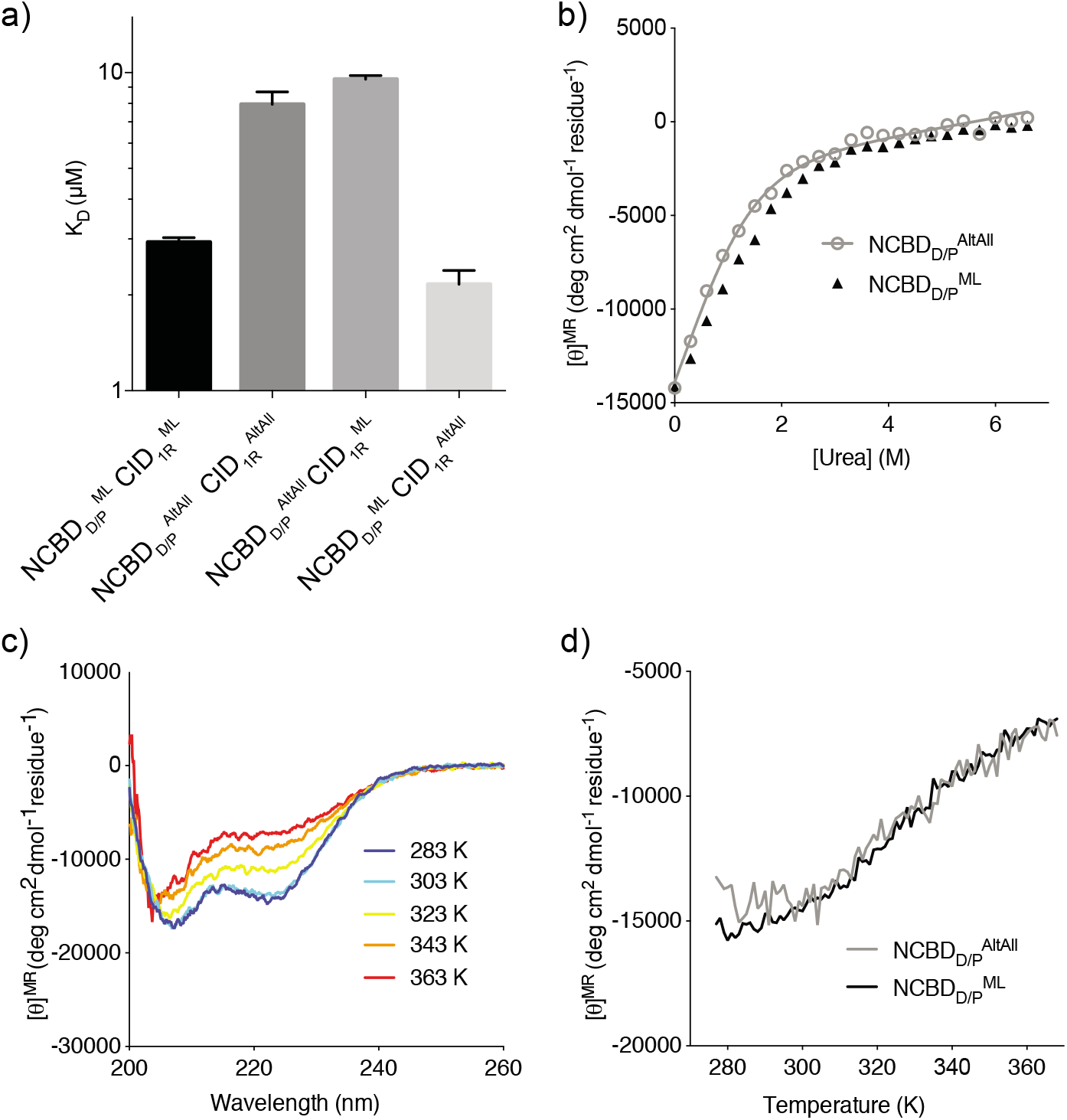
The biophysical properties of NCBD_D/P_^ML^ are robust to errors in the sequence. a) Binding affinities of NCBD_D/P_^ML^ or NCBD_D/P_^AltAlt^ and CID_1R_^ML^ or CID_1R_^AltAll^ measured with ITC. The buffer solution was 20 mM sodium phosphate pH 7.4, 150 mM NaCl. (b) Stability of NCBD_D/P_^AltAll^ measured by urea denaturation in 50 mM formate (pH 3.0), 150 mM NaCl. The CD signal at 222 nm was monitored and the data was fitted to a two-state function. A reliable [urea]_50%_ value could not be obtained, although qualitative comparison with NCBD_D/P_^ML^ indicates that the stability of the two variants is similar. (c-d) Temperature stability for NCBD_D/P_^AltAll^. The CD signal at 222 nm was monitored and spectra were taken at temperatures 283-363 K.

**Figure S3.**
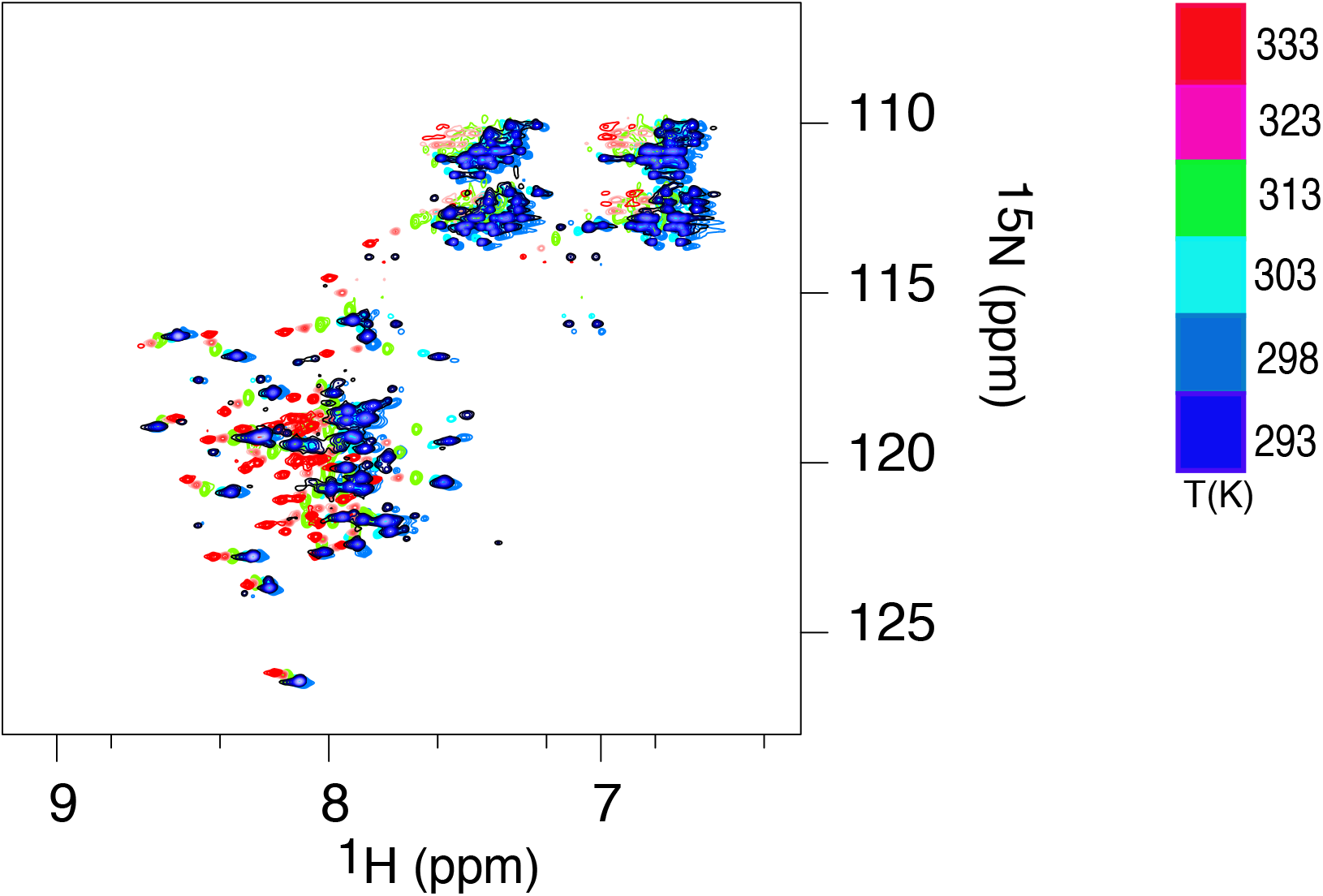
^15^N-^1^H HSQC spectra of NCBD_D/P_^ML^ at different temperatures. The experiments were performed from 293 K (blue) to 333 K (red).

**Figure S4.**
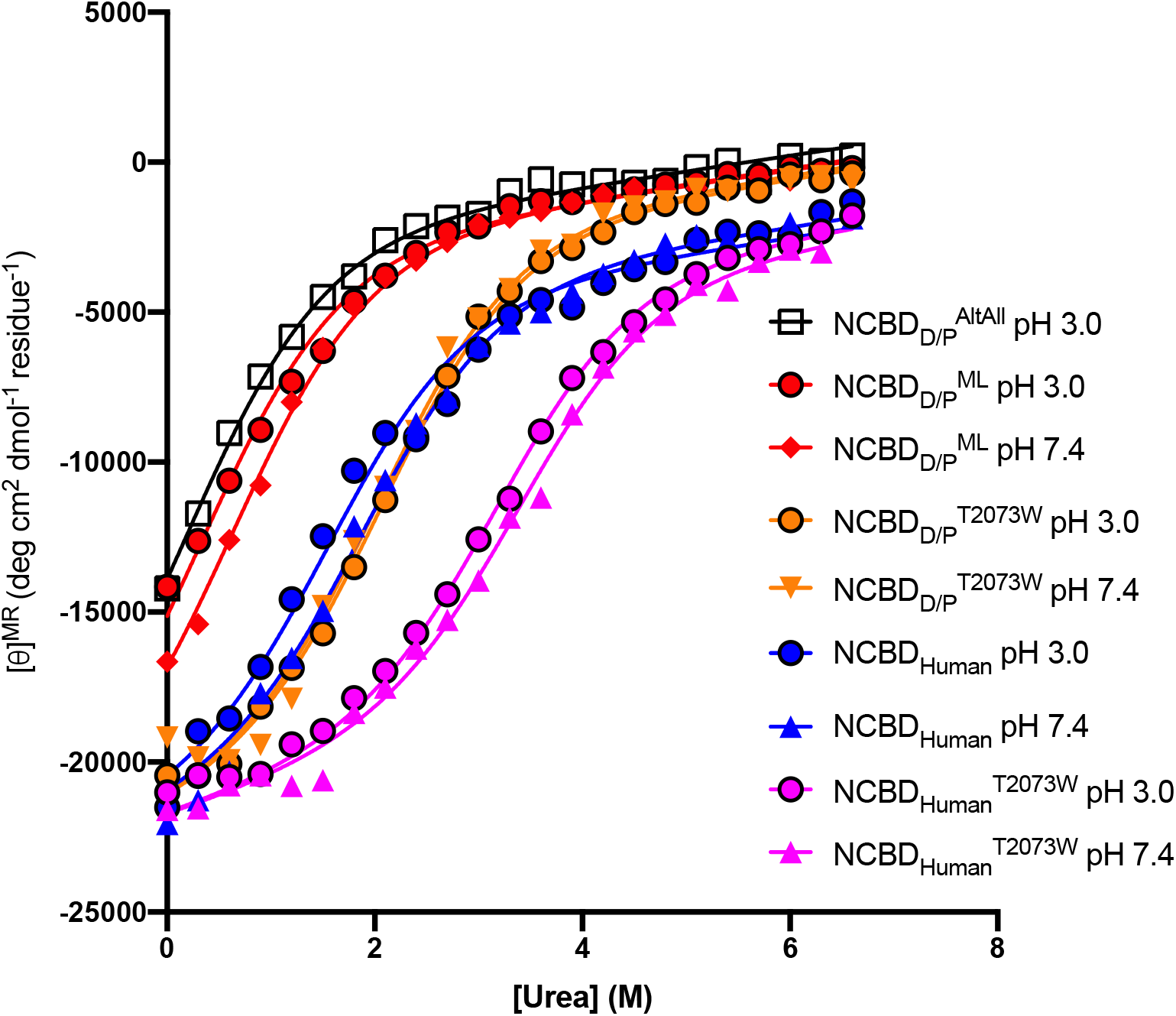
Equilibrium urea denaturation experiments. All data sets were fitted simultaneously to a two-state model in GraphPad Prism with shared parameters for the native state molar ellipticity, the denatured and native baseline slopes and the m_D-N_ value. Experiments were performed in either 50 mM potassium formate buffer (pH 3.0), 150 mM NaCl or 20 mM sodium phosphate buffer (pH 7.4), 150 mM NaCl. The shared m_D-N_ value was fitted as 0.74 ± 0.07 kcal mol^-1^M^-1^.

### Supplementary Tables

**Supplementary Table S1.** Stabilities of NCBD variants expressed as the urea concentration at 50% denaturation. The proteins were denatured with urea and the structural transition was monitored with CD at 222 nm at 277 K. The solutions were either 50 mM potassium formate buffer (pH 3.0), 150 mM NaCl or 20 mM sodium phosphate buffer (pH 7.4), 150 mM NaCl. The data were fitted to a two-state model in GraphPad Prism. The parameters for the signal of the native state, the denatured and native baseline slopes, and the m_D-N_ value were shared among the data sets in the curve fitting to allow fitting of the low stability NCBD_D/P_^ML^ and NCBD_D/P_^AltAll^ variants. The shared m_D-N_ value was fitted as 0.82 ± 0.025 kcal mol^-1^M^-1^. The errors are the standard error from the curve fitting and likely a low estimate of.

| Variant | pH | [Urea]<sub>50%</sub><br>(M) | $\Delta G_{D-N}$<br>(kcal/mol) |
| --- | --- | --- | --- |
| NCBD<sub>D/P</sub><sup>AltAll</sup> | 3.0 | $0.22 \pm 0.04$ | $0.18 \pm 0.03$ |
| NCBD<sub>D/P</sub><sup>ML</sup> | 3.0 | $0.38 \pm 0.04$ | $0.31 \pm 0.03$ |
| NCBD<sub>D/P</sub><sup>ML</sup> | 7.4 | $0.68 \pm 0.04$ | $0.55 \pm 0.04$ |
| NCBD<sub>D/P</sub><sup>T2073W</sup> | 3.0 | $2.13 \pm 0.04$ | $1.73 \pm 0.06$ |
| NCBD<sub>D/P</sub><sup>T2073W</sup> | 7.4 | $2.08 \pm 0.04$ | $1.70 \pm 0.06$ |
| NCBD<sub>Human</sub> | 3.0 | $1.61 \pm 0.04$ | $1.31 \pm 0.05$ |
| NCBD<sub>Human</sub> | 7.4 | $1.94 \pm 0.05$ | $1.58 \pm 0.06$ |
| NCBD<sub>Human</sub><sup>T2073W</sup> | 3.0 | $3.24 \pm 0.07$ | $2.64 \pm 0.10$ |
| NCBD<sub>Human</sub><sup>T2073W</sup> | 7.4 | $3.46 \pm 0.08$ | $2.82 \pm 0.11$ |

**Supplementary Table 2.** Rate constants and kinetic amplitudes of folding for different D/P NCBD Trp variants measured in temperature jump experiments. The experiments were conducted using 200 μM protein in 20 mM sodium phosphate (pH 7.4), 150 mM NaCl. The jump in temperature was from 277 to 285.5 K.

| **Variant** | **Amplitude** | **$k_{\text{obs}}$ ($\text{s}^{-1}$)** |
| --- | --- | --- |
| NCBD<sub>D/P</sub><sup>T2073W</sup> | $1.06 \pm 0.06$ | $2400 \pm 200$ |
| NCBD<sub>D/P</sub><sup>L2067W</sup> | $0.91 \pm 0.02$ | $1530 \pm 50$ |
| NCBD<sub>D/P</sub><sup>S2078W</sup> | $0.35 \pm 0.01$ | $700 \pm 60$ |
| NCBD<sub>D/P</sub><sup>H2107W</sup> | $0.47 \pm 0.02$ | $820 \pm 80$ |
| NCBD<sub>D/P</sub><sup>Q2108W</sup> | $0.73 \pm 0.02$ | $960 \pm 60$ |

**Supplementary Table 3.**

|  | NCBD<sub>D/P</sub><sup>ML</sup> | NCBD<sub>D/P</sub><sup>AltAlt</sup> |
| --- | --- | --- |
| **NMR distance and dihedral restraints** |  |  |
| **Distance restraints** |  |  |
| Total NOEs | 329 | 347 |
| Intra-residue | 113 | 145 |
| Sequential ($\vert i - j\vert = 1$) | 122 | 111 |
| Medium-range ($1 < \vert i - j\vert < 4$) | 76 | 76 |
| Long-range ($\vert i - j\vert > 5$) | 18 | 39 |
| Total dihedral angle restraints | 76 | 63 |
| $^3J_{\text{HN}\alpha}$ scalar couplings | 39 | 43 |
| $^{13}\text{C}^\alpha$ chemical shifts | 100 | 100 |
| **Structure statistics** |  |  |
| Average CYANA target function value ($\text{\AA}^2$) | $2.5 \pm 0.2$ | $2.8 \pm 0.3$ |
| Violations |  |  |
| Distance constraints ($>0.5 \text{ \AA}$) | 0 | 0 |
| Dihedral angle constraints ($>5^\circ$) | 0 | 0 |
| Deviations from idealized geometry |  |  |
| Bond lengths ($\text{\AA}$) | 0 | 0 |
| Bond angles ($^\circ$) | 0 | 0 |
| Impropers ($^\circ$) | 0 | 0 |
| Average pairwise r.m.s. deviation ($\text{\AA}$) | | |
| Backbone residues | $2.2 \pm 1.0$ | $0.5 \pm 0.2$ |
| Heavy atoms residues | $2.7 \pm 1.0$ | $1.0 \pm 0.3$ |

## References

1. S. Gianni, M. I. Freiberger, P. Jemth, D. U. Ferreiro, P. G. Wolynes, M. Fuxreiter, Fuzziness and Frustration in the Energy Landscape of Protein Folding, Function, and Assembly. Acc Chem Res. 54, 1251–1259 (2021).

2. O. F. Lange, N.-A. Lakomek, C. Farès, G. F. Schröder, K. F. A. Walter, S. Becker, J. Meiler, H. Grubmüller, C. Griesinger, B. L. de Groot, Recognition dynamics up to microseconds revealed from an RDC-derived ubiquitin ensemble in solution. Science. 320, 1471–1475 (2008).

3. S. J. Wodak, E. Paci, N. V. Dokholyan, I. N. Berezovsky, A. Horovitz, J. Li, V. J. Hilser, I. Bahar, J. Karanicolas, G. Stock, P. Hamm, R. H. Stote, J. Eberhardt, Y. Chebaro, A. Dejaegere, M. Cecchini, J.-P. Changeux, P. G. Bolhuis, J. Vreede, P. Faccioli, S. Orioli, R. Ravasio, L. Yan, C. Brito, M. Wyart, P. Gkeka, I. Rivalta, G. Palermo, J. A. McCammon, J. Panecka-Hofman, R. C. Wade, A. Di Pizio, M. Y. Niv, R. Nussinov, C.-J. Tsai, H. Jang, D. Padhorny, D. Kozakov, T. McLeish, Allostery in Its Many Disguises: From Theory to Applications. Structure. 27, 566–578 (2019).

4. H. J. Dyson, P. E. Wright, Role of Intrinsic Protein Disorder in the Function and Interactions of the Transcriptional Coactivators CREB-binding Protein (CBP) and p300. J. Biol. Chem. 291, 6714–22 (2016).

5. C. H. Lin, B. J. Hare, G. Wagner, S. C. Harrison, T. Maniatis, E. Fraenkel, A small domain of CBP/p300 binds diverse proteins: solution structure and functional studies. Mol Cell. 8, 581–590 (2001).

6. S. J. Demarest, M. Martinez-Yamout, J. Chung, H. Chen, W. Xu, H. J. Dyson, R. M. Evans, P. E. Wright, Mutual synergistic folding in recruitment of CBP/p300 by p160 nuclear receptor coactivators. Nature. 415, 549–553 (2002).

7. M.-O. Ebert, S.-H. Bae, H. J. Dyson, P. E. Wright, NMR relaxation study of the complex formed between CBP and the activation domain of the nuclear hormone receptor coactivator ACTR. Biochemistry. 47, 1299–308 (2008).

8. M. Kjaergaard, K. Teilum, F. M. Poulsen, Conformational selection in the molten globule state of the nuclear coactivator binding domain of CBP. Proc. Natl. Acad. Sci. U.S.A. 107, 12535–12540 (2010).

9. M. Kjaergaard, F. M. Poulsen, K. Teilum, Is a malleable protein necessarily highly dynamic? The hydrophobic core of the nuclear coactivator binding domain is well ordered. Biophys J. 102, 1627–1635 (2012).

10. M. Kjaergaard, L. Andersen, L. D. Nielsen, K. Teilum, A folded excited state of ligand-free nuclear coactivator binding domain (NCBD) underlies plasticity in ligand recognition. Biochemistry. 52, 1686–1693 (2013).

11. E. Papaleo, C. Camilloni, K. Teilum, M. Vendruscolo, K. Lindorff-Larsen, Molecular dynamics ensemble refinement of the heterogeneous native state of NCBD using chemical shifts and NOEs. PeerJ. 6, e5125 (2018).

12. G. Hultqvist, E. Åberg, C. Camilloni, G. N. Sundell, E. Andersson, J. Dogan, C. N. Chi, M. Vendruscolo, P. Jemth, Emergence and evolution of an interaction between intrinsically disordered proteins. Elife. 6, e16059 (2017).

13. P. Jemth, E. Karlsson, B. Vögeli, B. Guzovsky, E. Andersson, G. Hultqvist, J. Dogan, P. Güntert, R. Riek, C. N. Chi, Structure and dynamics conspire in the evolution of affinity between intrinsically disordered proteins. Sci. Adv. 4, eaau4130 (2018).

14. G. N. Eick, J. T. Bridgham, D. P. Anderson, M. J. Harms, J. W. Thornton, Robustness of Reconstructed Ancestral Protein Functions to Statistical Uncertainty. Mol Biol Evol. msw223 (2016).

15. K. M. Hart, M. J. Harms, B. H. Schmidt, C. Elya, J. W. Thornton, S. Marqusee, Thermodynamic system drift in protein evolution. PLoS Biol. 12, e1001994 (2014).

16. L. Laursen, J. Čalyševa, T. J. Gibson, P. Jemth, Divergent Evolution of a Protein-Protein Interaction Revealed through Ancestral Sequence Reconstruction and Resurrection. Mol Biol Evol. 38, 152–167 (2021).

17. P. D. Williams, D. D. Pollock, B. P. Blackburne, R. A. Goldstein, Assessing the accuracy of ancestral protein reconstruction methods. PLoS Comput Biol. 2, e69 (2006).

18. V. A. Risso, J. A. Gavira, E. A. Gaucher, J. M. Sanchez-Ruiz, Phenotypic comparisons of consensus variants versus laboratory resurrections of Precambrian proteins. Proteins. 82, 887–896 (2014).

19. M. L. Romero-Romero, V. A. Risso, S. Martinez-Rodriguez, B. Ibarra-Molero, J. M. Sanchez-Ruiz, Engineering ancestral protein hyperstability. Biochem J. 473, 3611–3620 (2016).

20. D. L. Trudeau, M. Kaltenbach, D. S. Tawfik, On the Potential Origins of the High Stability of Reconstructed Ancestral Proteins. Mol Biol Evol. 33, 2633–2641 (2016).

21. L. C. Wheeler, S. A. Lim, S. Marqusee, M. J. Harms, The thermostability and specificity of ancient proteins. Curr Opin Struct Biol. 38, 37–43 (2016).

22. T. W. Hearing, T. H. P. Harvey, M. Williams, M. J. Leng, A. L. Lamb, P. R. Wilby, S. E. Gabbott, A. Pohl, Y. Donnadieu, An early Cambrian greenhouse climate. Sci Adv. 4, eaar5690 (2018).

23. T. Wotte, C. B. Skovsted, M. J. Whitehouse, A. Kouchinsky, Isotopic evidence for temperate oceans during the Cambrian Explosion. Sci Rep. 9, 6330 (2019).

24. J. Dogan, A. Toto, E. Andersson, S. Gianni, P. Jemth, Activation Barrier-Limited Folding and Conformational Sampling of a Dynamic Protein Domain. Biochemistry. 55, 5289–5295 (2016).

25. S. J. Demarest, S. Deechongkit, H. J. Dyson, R. M. Evans, P. E. Wright, Packing, specificity, and mutability at the binding interface between the p160 coactivator and CREB-binding protein. Protein Sci. 13, 203–210 (2004).

26. J. Xu, Q. Li, Review of the in vivo functions of the p160 steroid receptor coactivator family. Mol. Endocrinol. 17, 1681–1692 (2003).

27. J. A. Livengood, K. E. S. Scoggin, K. Van Orden, S. J. McBryant, R. S. Edayathumangalam, P. J. Laybourn, J. K. Nyborg, p53 Transcriptional activity is mediated through the SRC1-interacting domain of CBP/p300. J Biol Chem. 277, 9054–9061 (2002).

28. G. Jayaraman, R. Srinivas, C. Duggan, E. Ferreira, S. Swaminathan, K. Somasundaram, J. Williams, C. Hauser, M. Kurkinen, R. Dhar, S. Weitzman, G. Buttice, B. Thimmapaya, p300/cAMP-responsive element-binding protein interactions with ets-1 and ets-2 in the transcriptional activation of the human stromelysin promoter. J Biol Chem. 274, 17342–17352 (1999).

29. S. Matsuda, J. C. Harries, M. Viskaduraki, P. J. F. Troke, K. B. Kindle, C. Ryan, D. M. Heery, A Conserved alpha-helical motif mediates the binding of diverse nuclear proteins to the SRC1 interaction domain of CBP. J Biol Chem. 279, 14055–14064 (2004).

30. K. E. Scoggin, A. Ulloa, J. K. Nyborg, The oncoprotein Tax binds the SRC-1-interacting domain of CBP/p300 to mediate transcriptional activation. Mol Cell Biol. 21, 5520–5530 (2001).

31. B. Y. Qin, C. Liu, H. Srinath, S. S. Lam, J. J. Correia, R. Derynck, K. Lin, Crystal structure of IRF-3 in complex with CBP. Structure. 13, 1269–1277 (2005).

32. E. Karlsson, A. Lindberg, E. Andersson, P. Jemth, High affinity between CREBBP/p300 and NCOA evolved in vertebrates. Protein Sci. 29, 1687–1691 (2020).

33. E. Karlsson, C. Paissoni, A. M. Erkelens, Z. A. Tehranizadeh, F. A. Sorgenfrei, E. Andersson, W. Ye, C. Camilloni, P. Jemth, Mapping the transition state for a binding reaction between ancient intrinsically disordered proteins. J Biol Chem. 295, 17698–17712 (2020).

34. P. E. Wright, H. J. Dyson, Intrinsically disordered proteins in cellular signalling and regulation. Nat Rev Mol Cell Biol. 16, 18–29 (2015).

35. A. Toto, C. Camilloni, R. Giri, M. Brunori, M. Vendruscolo, S. Gianni, T. A, C. C, G. R, B. M, V. M, G. S., Molecular Recognition by Templated Folding of an Intrinsically Disordered Protein. Sci. Rep. 6, 21994 (2016).

36. A. Toto, F. Malagrinò, L. Visconti, F. Troilo, L. Pagano, M. Brunori, P. Jemth, S. Gianni, Templated folding of intrinsically disordered proteins. J. Biol. Chem. 295, 6586–6593 (2020).

37. D. Wu, H.-X. Zhou, W. D, Z. HX., Designed Mutations Alter the Binding Pathways of an Intrinsically Disordered Protein. Sci. Rep. 9, 6172 (2019).

38. P. Jemth, X. Mu, Å. Engström, J. Dogan, A frustrated binding interface for intrinsically disordered proteins. J. Biol. Chem. 289, 5528–5533 (2014).

39. E. Karlsson, E. Andersson, J. Dogan, S. Gianni, P. Jemth, C. Camilloni, A structurally heterogeneous transition state underlies coupled binding and folding of disordered proteins. J. Biol. Chem. 294, 1230–1239 (2019).

40. C. Chothia, A. M. Lesk, The relation between the divergence of sequence and structure in proteins. EMBO J. 5, 823–826 (1986).

41. M. Oliveberg, P. G. Wolynes, The experimental survey of protein-folding energy landscapes. Q. Rev. Biophys. 38, 245–288 (2005).

42. A. Fersht, Structure and mechanism in protein science: a guide to enzyme catalysis and protein folding. (Macmillan, 1999).

43. N.-A. Lakomek, J. Ying, A. Bax, Measurement of 15N relaxation rates in perdeuterated proteins by TROSY-based methods. J Biomol NMR. 53, 209–221 (2012).

44. F. Delaglio, S. Grzesiek, G. W. Vuister, G. Zhu, J. Pfeifer, A. Bax, NMRPipe: a multidimensional spectral processing system based on UNIX pipes. J Biomol NMR. 6, 277–293 (1995).

45. W. F. Vranken, W. Boucher, T. J. Stevens, R. H. Fogh, A. Pajon, M. Llinas, E. L. Ulrich, J. L. Markley, J. Ionides, E. D. Laue, The CCPN data model for NMR spectroscopy: development of a software pipeline. Proteins. 59, 687–696 (2005).

46. P. Güntert, C. Mumenthaler, K. Wüthrich, Torsion angle dynamics for NMR structure calculation with the new program Dyana. J. Mol. Biol. 273, 283–298 (1997).

